# An intranasal ASO therapeutic targeting SARS-CoV-2

**DOI:** 10.1101/2021.05.17.444397

**Authors:** Chi Zhu, Justin Y. Lee, Jia Z. Woo, Lei Xu, Xammy Nguyenla, Livia H. Yamashiro, Fei Ji, Scott B. Biering, Erik Van Dis, Federico Gonzalez, Douglas Fox, Arjun Rustagi, Benjamin A. Pinsky, Catherine A. Blish, Charles Chiu, Eva Harris, Ruslan I. Sadreyev, Sarah Stanley, Sakari Kauppinen, Silvi Rouskin, Anders M. Näär

**Author notes:** Corresponding author, Correspondence to Anders M. Näär.

## Abstract

The COVID-19 pandemic is exacting an increasing toll worldwide, with new SARS-CoV-2 variants emerging that exhibit higher infectivity rates and that may partially evade vaccine and antibody immunity^1^. Rapid deployment of non-invasive therapeutic avenues capable of preventing infection by all SARS-CoV-2 variants could complement current vaccination efforts and help turn the tide on the COVID-19 pandemic^2^. Here, we describe a novel therapeutic strategy targeting the SARS-CoV-2 RNA using locked nucleic acid antisense oligonucleotides (LNA ASOs). We identified an LNA ASO binding to the 5’ leader sequence of SARS-CoV-2 ORF1a/b that disrupts a highly conserved stem-loop structure with nanomolar efficacy in preventing viral replication in human cells. Daily intranasal administration of this LNA ASO in the K18-hACE2 humanized COVID-19 mouse model potently (98-99%) suppressed viral replication in the lungs of infected mice, revealing strong prophylactic and treatment effects. We found that the LNA ASO also represses viral infection in golden Syrian hamsters, and is highly efficacious in countering all SARS-CoV-2 “variants of concern” tested *in vitro* and *in vivo*, including B.1.427, B.1.1.7, and B.1.351 variants^3^. Hence, inhaled LNA ASOs targeting SARS-CoV-2 represents a promising therapeutic approach to reduce transmission of variants partially resistant to vaccines and monoclonal antibodies, and could be deployed intranasally for prophylaxis or via lung delivery by nebulizer to decrease severity of COVID-19 in infected individuals. LNA ASOs are chemically stable and can be flexibly modified to target different viral RNA sequences^4^, and they may have particular impact in areas where vaccine distribution is a challenge, and could be stockpiled for future coronavirus pandemics.

## Main

The global coronavirus disease 19 (COVID-19) pandemic is caused by the highly pathogenic novel human SARS-coronavirus 2 (SARS-CoV-2)^5^. By contrast with previous outbreaks of related beta-coronaviruses (severe acute respiratory syndrome coronavirus (SARS-CoV-1) and Middle East respiratory syndrome conronavirus (MERS-CoV)), SARS-CoV-2 infection causes lower mortality rate, however, the virus has a higher human-to-human transmission rate^6^, facilitating rapid spread across the world. As of May 2021, there are more than 150 million confirmed positive cases and in excess of 3 million reported deaths (WHO; www.who.int). Although vaccines are markedly slowing the increase in positive cases and deaths in a few developed countries, most nations, especially in the developing world, do not have readily accessible vaccines^7^, and there is aversion among segments of the global population to vaccination^7^. Moreover, mutated variant strains of SARS-CoV-2 that evade immunity in response to previous infection or vaccination are rapidly emerging, and are causing new local outbreaks, frequently amongst younger individuals and with more severe disease^1,8^. Indeed, more than 80 variants have been identified to date, all with mutations in the Spike protein^9^, which SARS-CoV-2 utilizes for binding to the host cell-surface receptor angiotensin-converting enzyme 2 (ACE2) during host cell entry^10^. Several “variants of concern”, which harbor mutations in Spike that facilitate immune evasion and, in some cases, partial vaccine and monoclonal antibody therapeutics resistance, are spreading rapidly world-wide. These include B.1.1.7 (UK variant), B.1.351 (South African/SA variant), P.1 (Brazilian variant), B.1.427/B.1.429 (CA variants), and B.1.617 (Indian variant), which are all more contagious and may in some cases lead to more severe disease^11^. The continuous evolution of the virus thus poses a daunting challenge to achieving “herd immunity”. Traditional drug screening and vaccine development is time intensive, and may not be able to match the speed of emerging drug- or vaccine-resistant SARS-CoV-2 strains. There is clearly an urgent need for alternative approaches, including the rapid development of therapeutics that are active against all variants of concern.

To address these challenges, we developed a novel strategy to inhibit the replication of SARS-CoV-2 using antisense oligonucleotides (ASOs) targeting viral RNAs. ASOs, which rely on Watson-Crick base-pairing to target specific complementary RNA sequences, can be quickly designed to target any viral or host RNA sequence, including non-coding structural elements that may be important for viral replication, and may recruit RNase H for cleavage (gapmers) or act through steric hindrance (mixmers)^4^. ASOs are typically well tolerated, and a number of ASO therapeutics have been approved for clinical use^12^. Additionally, ASO manufacturing is well established and can be readily scaled-up. We have employed chemically modified gapmer and mixmer ASOs containing interspersed locked nucleic acid nucleotide bases (LNAs) and DNA nucleotides linked by phosphorothioate (PS) bonds. The introduced chemical modifications confer increased affinity, stability and improved pharmacokinetic/pharmacodynamic properties^13,14,15^.

SARS-CoV-2 is a compact (30 kilobases) positive-sense single-stranded RNA virus, with a 5’ untranslated region (UTR), the ORF1a/b RNA encoding non-structural viral proteins, and a 3’ segment encoding the structural RNAs, such as the Spike protein that binds to the ACE2 receptor on host cells, and the nucleocapsid N protein involved in virion assembly, and a 3’UTR^16^. The 5’ UTR, a non-coding segment consisting of multiple highly conserved stem-loop and more complex secondary structures, is functionally critical for viral translation and replication by affording protection from host cell antiviral defenses and through selective promotion of viral transcript translation over those of the host cell, at least in part through the recruitment of the viral non-strctural protein 1 (Nsp1)^17^. The 5’ UTR begins with a short 5’ leader sequence (nucleotides 1-69), which is added via discontinuous transcription to the 5’ end of all sub-genomic RNA transcripts encoding the viral structural proteins, and regulates their translation as well as translation of ORF1a/b from full-length genomic RNA^18^. The ORF1a/b also contains a structured and highly conserved frameshift stimulation element (FSE) near its center that controls a shift in the protein translation reading frame by one nucleotide of ORF1a/b genes 3’ to the FSE. The FSE and accurate frame shifting is crucial for the expression of ORF1b, which encodes five non-structural proteins including an RNA-dependent RNA polymerase (RdRP) essential for SARS-CoV-2 genome replication^19^.

We designed multiple LNA ASOs targeting the 5’ leader sequence, downstream sequences in the 5’ untranslated region (UTR) of ORF1a/b, and the ORF1a/b FSE of SARS-CoV-2 (Fig. 1a and Extended Data Table 1). Additionally, given the importance of the Spike protein in host cell entry for SARS-CoV-2 infection, LNA ASOs targeting the Spike coding sequence were also tested (Fig. 1a and Extended Data Table 1). To evaluate the viral repressive effect of LNA ASOs, the initial screening was carried out in Huh-7 human hepatoma cells, which exhibit excellent transfection efficiency and are readily infected by SARS-CoV-2. Cells transfected with LNA ASOs were infected with SARS-CoV-2 and both cells and medium were collected at 48h post-infection (hpi) for RNA extraction and infectious viral particle determination, respectively. The viral titer was measured by detecting the expression level/copy number of *Nucleocapsid* (*N*) and *Spike* (*S*) using reverse transcription (RT)-quantitative PCR (qPCR). The screening results showed that the treatment with certain LNA ASOs lead to a dramatic decrease of *N* and *S* expression (Fig. 1b and 1c, Extended Data Fig. 1a and 1b).

**Figure 1.**
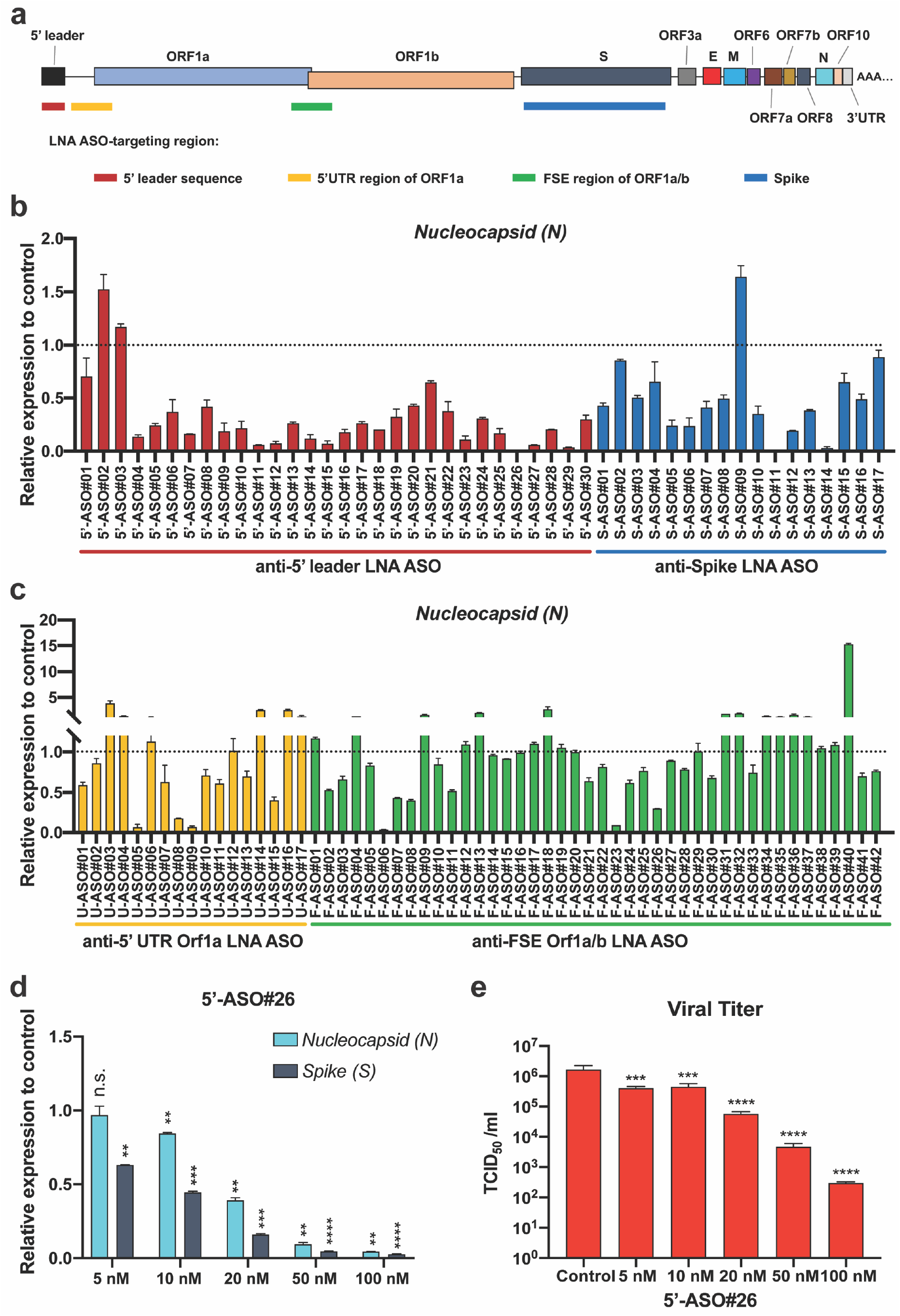
*In vitro* screening of LNA ASOs targeting SARS-CoV-2. a) Schematic representation of the genome structure and the regions targeted by LNA ASOs. b) and c) The SARS-CoV-2 WA1 strain-infected Huh-7 cells treated with LNA ASO (100 nM) and cell culture medium were collected at 48 hpi. Viral RNA levels were analyzed by RT-qPCR for LNA ASO screening. Each LNA ASO was tested in duplicate. d) Dose-dependent effects of 5’-ASO#26 were evaluated in infected Huh-7 cells with increasing doses of the LNA ASO (as indicated) by RT-qPCR. e) The infectious virus was measured by TCID_50_ assay. The infected Huh-7 cells with different doses of LNA ASO treatment were collected at 48 hpi. For d) and e), one-way ANOVA with Dunnett’s test was used to determine significance (** *P* < 0.01, *** *P* < 0.001, **** *P* < 0.0001, ns, not significant).

Interestingly, we found that LNA ASOs targeting the 5’ leader region of SARS-CoV-2 were particularly effective in suppressing viral RNA levels in infected cells (Fig. 1b and Extended Data Fig. 1a). This is consistent with the fact that the 5’ leader sequence is present in all viral RNA transcripts and is required for viral replication. The most potent LNA ASO targeting the 5’ leader, 5’-ASO#26, was selected for further investigation of its viral repression capability. The repressive effect of 5’-ASO#26 was demonstrated in a dose-dependent manner by measuring the expression level of viral RNAs (Fig. 1d) and by directly determining the viral titer of infectious particles with the Fifty-percent Tissue Culture Infectious Dose (TCID_50_) assay in Vero E6 African green monkey kidney epithelial cells (Fig. 1e).

Although the 5’ UTR nucleotide sequences are somewhat divergent amongst the coronavirus family, the secondary structure of the 5’ UTR is highly conserved^20^, and it has been shown that two stem-loop structures, SL1 and SL2, are formed by the 5’ leader sequence^21^. Since the complementary sequence of 5’-ASO#26 aligns along the 3’ portion of SL1 (marked in pink frame) (Fig. 2c), we hypothesized that the viral repressive effect of 5’-ASO#26 may be in part due to its ability to disrupt the secondary structure of the 5’ leader sequence upon binding to the viral genomic or sub-genomic RNAs, interfering with the formation of the SL1 stem-loop structure. To test if 5’-ASO#26 can disrupt the SL1 structure, we carried out *in vitro* transcription of the SARS-CoV-2 5’ UTR sequence, added 5’-ASO#26 and treated samples with dimethyl sulfate (DMS). DMS is able to specifically and rapidly methylate unpaired adenines (A) and cytosines (C) within single-stranded sequences and not those that are complexed as RNA secondary structure or to the LNA ASO, allowing for the unpaired A or C nucleotides to be detected by DMS-MaPseq^22^. Our results strongly indicated that the secondary structure of SL1 was indeed interrupted by 5’-ASO#26 in a dose-dependent manner (Fig. 2a). We also performed DMS-MaPseq with SARS-CoV-2-infected Huh-7 cells in the presence or absence of 5’-ASO#26, and monitored secondary structure changes of SL1 at 6 hpi and 12 hpi. Similar to the findings with the *in vitro* assay, the result from infected cells confirmed the disruption of the secondary structure of SL1 due to binding of 5’-ASO#26 (Fig. 2b).

**Figure 2.**
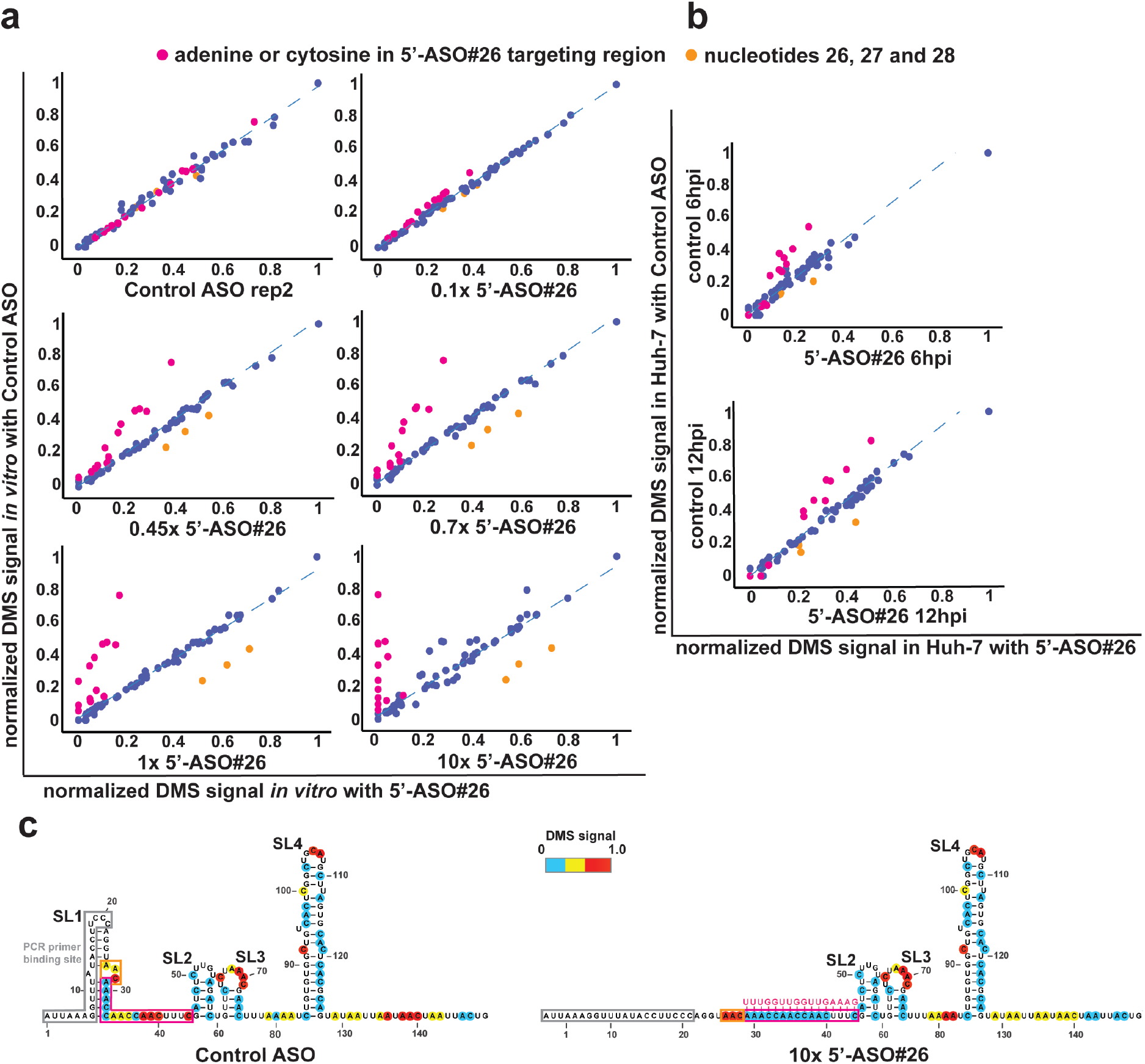
5’-ASO#26 disrupts the stem-loop structure of SL1. a) Scatter plots comparing 5’-leader structures *in vitro* in different conditions. Comparison of DMS structure signals between the control LNA ASO replicates (left top), and control verses titration with 5’-ASO#26 at 0.1x, 0.45x, 0.7x, 1x and 10x molar ratio of 5’-ASO#26 to the 5’-leader RNA. b) Scatter plots comparing 5’-leader structures in infected Huh-7 cells at different time points. Comparison of DMS signals between samples transfected with control LNA ASO and with 5’-ASO#26 at 6 and 12 dpi. For a) and b) blue dotted line indicates the fitted regression line. Every dot represents an adenine or cytosine in the 5’leader. Pink dots represent adenines or cytosines in the 5’-ASO#26 targeting region. Orange dots represent nucleotides 26, 27 and 28 of the 5’ leader. c) Structural model of SL1 predicted from the DMS-MaPseq of *in vitro*-transcribed 5’leader RNA with the addition of control LNA ASO (left) and 5’-ASO#26 (right). Nucleotides are color-coded by normalized DMS signal. The 5’-ASO#26-binding site is highlighted with a pink frame; nucleotides 26, 27 and 28 are highlighted with an orange frame; PCR primer binding site, where DMS information is unavailable, is highlighted with a grey frame.

To evaluate the effects of 5’-ASO#26 *in vivo*, we employed humanized transgenic K18-hACE2 mice, which are expressing human ACE2 allowing SARS-CoV-2 cell entry and infectious spread^23^. K18-hACE2 mice were inoculated with 1×10^4^ TCID_50_ units of the USA-WA1/2020 SARS-CoV-2 strain via intranasal administration. No significant weight loss was observed at 4 days post-infection (dpi) (Extended Data Fig. 2a), consistent with previous studies indicating that weight loss starts from 5 dpi^23^. Mice were treated with 5’-ASO#26 (400 μg) once-daily via intranasal administration in saline, from 3 days before infection until 3 dpi (Fig. 3a). High levels of infectious SARS-CoV-2 viral particles were detected in the lungs of the saline-treated control group (Saline), whereas a remarkable decrease (>80-fold) in the viral load in lungs was observed in the LNA ASO-treated group (Fig. 3b). As expected, the levels of viral *N* and *S* RNA were also potently (98-99%) repressed after LNA ASO treatment, as measured by RT-qPCR (Fig. 3c). Previous studies have shown that golden Syrian hamsters can be infected with SARS-CoV-2^24^. We inoculated hamsters with 10 TCID_50_ units of the USA-WA1/2020 SARS-CoV-2 strain and treated them with 5’-ASO#26 (600 μg) following the same schedule as for the mouse studies (Fig. 3a). About 10-fold decreased viral load in lung was observed in the LNA ASO-treated group and with no significant weight change (Extended Data Fig. 2d, 2e). Considering that the dose was approximately 2.5-fold lower per kg, the viral replication was still efficiently repressed in hamsters. To further investigate the physiological effects of LNA ASO in mice, histological analyses were carried out with control and LNA ASO-treated lung tissue after SARS-CoV-2 infection. The analyses revealed clear repression of N expression with LNA ASO treatment (Fig. 3d). Meanwhile, we did not observe significant histological changes by H&E staining in either Saline or LNA ASO-treated mice, which is consistent with previous studies demonstrating that significant inflammation and severe alveolar wall thickening could only be observed at 7 dpi^23^. However, we did notice that the localized signal of N correlated with local enrichment of CD3 (T cell marker), B220 (B cell marker) by immunohistochemistry, and moderate thickening of the alveolar wall (by H&E staining) in Saline-treated mice, but not in LNA ASO-treated mice (Extended Data Fig. 2c). The viral repressive effect of the LNA ASO was also examined by RNA-seq of lung tissues from mice with saline or LNA ASO treatment. Consistent with results from the TCID_50_ assay, viral reads were dramatically decreased in LNA ASO-treated mice when compared with that of the Saline control (Extended Data Fig. 3a). To evaluate the *in vivo* effect of 5’-ASO#26, previous RNA-seq data from infected K18-ACE2 mice^23^ was used to define up- and down-regulated genes in response to SARS-CoV-2 infection at 4 dpi (Extended Data Fig. 3e). As expected, expression profile changes induced by infection were markedly rescued by LNA ASO treatment (Fig. 3e and Extended Data Fig. 3b). Gene set enrichment analysis (GSEA) revealed strong enrichment of gene involved in type I and II IFN signaling in infected mice treated with Saline (Fig. 3f and Extended Data Fig. 3c), which is consistent with previous studies showing induction of anti-viral defenses^23^. Interestingly, the cholesterol homeostasis gene pathway was enriched in LNA ASO-treated mice (Fig. 3f and Extended Data Fig. 3d). It has been reported that the cholesterol biosynthesis pathway is necessary for SARS-CoV-2 infection^25,26^ and decreased levels of total cholesterol, high-density lipoprotein (HDL), and low-density lipoprotein (LDL) were reported in COVID-19 patients^27^, indicating a comprehensive role of intracellular and extracellular cholesterol during SARS-CoV-2 infection. The restored cholesterol homeostasis in the lungs of LNA ASO-treated mice further indicates the efficacy of the LNA ASO treatment.

**Figure 3.**
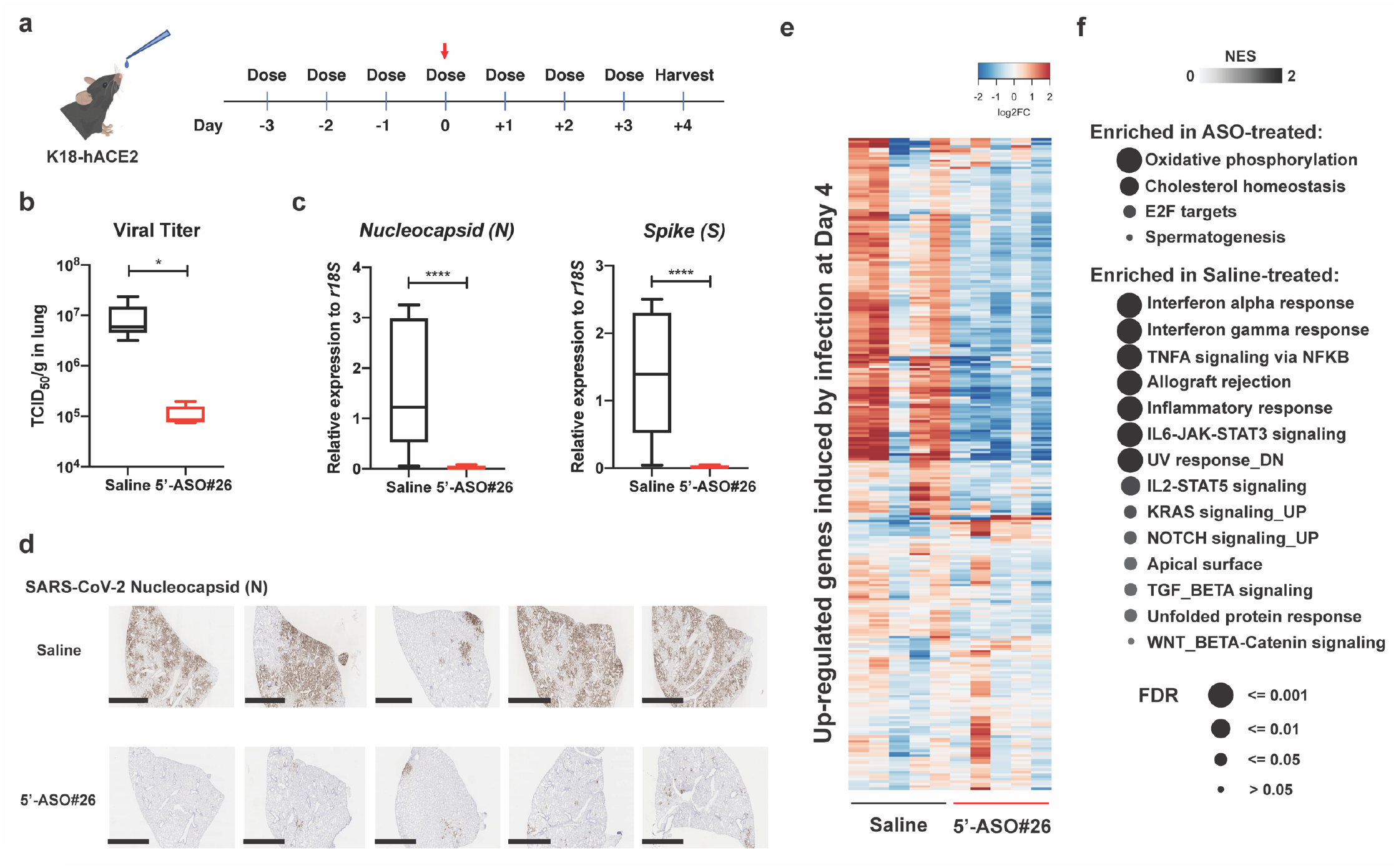
Once-daily treatment of 5’-ASO#26 represses the generation of infectious virus in mice. a) Treatment course of once-daily 5’-ASO#26 administration. The red arrow indicates the inoculation of 1×10^4^ TCID_50_ SARS-CoV-2 intranasally in mice. Mice were treated once-daily with saline (vehicle) or 400 μg LNA ASO (dissolved in saline) and hamsters were treated once-daily with saline (vehicle) or 600 μg LNA ASO (dissolved in saline). Treatment on the day of infection was carried out at 6 hpi. b) The viral burden in lungs of mice treated with Saline (N=5) or LNA ASO (N =5) was measured by TCID_50_ assay using lung homogenates. c) The levels of viral RNAs encoding Nucleocapsid protein (N) and Spike (S) were measured in mouse lungs by RT-qPCR. For b) and c), center line, median; box limits, upper and lower quartiles; plot limits, maximum and minimum in the boxplot. Student *t*-test was used to determine significance (* *P* < 0.05, **** *P* < 0.0001). d) IHC staining of SARS-CoV-2 N in all infected K18-hACE2 mice with or without LNA ASO treatment. Scale bar = 2 mm. e) Expression changes of SARS-CoV-2 infection-upregulated genes in Saline- and LNA ASO-treated groups. Columns represent samples and rows represent genes. Colors indicate expression levels (log2 RPKM) relative to average expression across all samples. f) Gene set enrichment analysis (GSEA) of Hallmark gene sets enriched in lungs of Saline- or LNA ASO-treated mice. Terms were ranked by the false discovery rate (q value).

To further evaluate the effect of 5’-ASO#26, we first confirmed that 5’-ASO#26 repressed viral replication *in vivo* in a dose-dependent manner (Extended Data Fig. 2b). To explore the optimal treatment time course, we assessed the efficacy of 5’-ASO#26 viral repression in pre-infection and post-infection treatment regimens (Fig. 4a). We found that the strongest viral repressive effect was observed in the Prophylactic #2 group (Fig. 4a), in which the mice were treated with LNA ASO for four days, followed by infection 24 hrs later. Viral repression was also validated by staining of N in mouse lung tissues (Extended Data Fig. 4a). Notably, a prophylactic effect was still observed one week after LNA ASO treatment (Prophylactic #1, Fig. 4a). Starting LNA ASO treatment at 6 hpi also revealed a moderate repressive effect, whereas beginning treatment from 1 dpi showed no repressive effect in mice (Treatment #3 and #4, Fig. 4a). These results show that 5’-ASO#26 exhibits potent effects as a prophylactic therapeutic. The diminished ability of late post-infection treatment of 5’-ASO#26 to inhibit viral replication may be a result of rapidly accumulating sub-genomic viral RNAs saturating the amount of administered LNA ASO in the lung due to the very large dose of virus used to inoculate the mice. Considering that we directly administered the naked 5’-ASO#26 intranasally in saline, we believe that additional modifications of the LNA ASO (such as lipid-conjugation^28,29,30^) may further promote cellular uptake of LNA ASO in lung, improving the effect of post-infection LNA ASO treatment. Direct delivery of the LNA ASO to the lung via nebulizer may also improve post-infection therapeutic effect.

**Figure 4.**
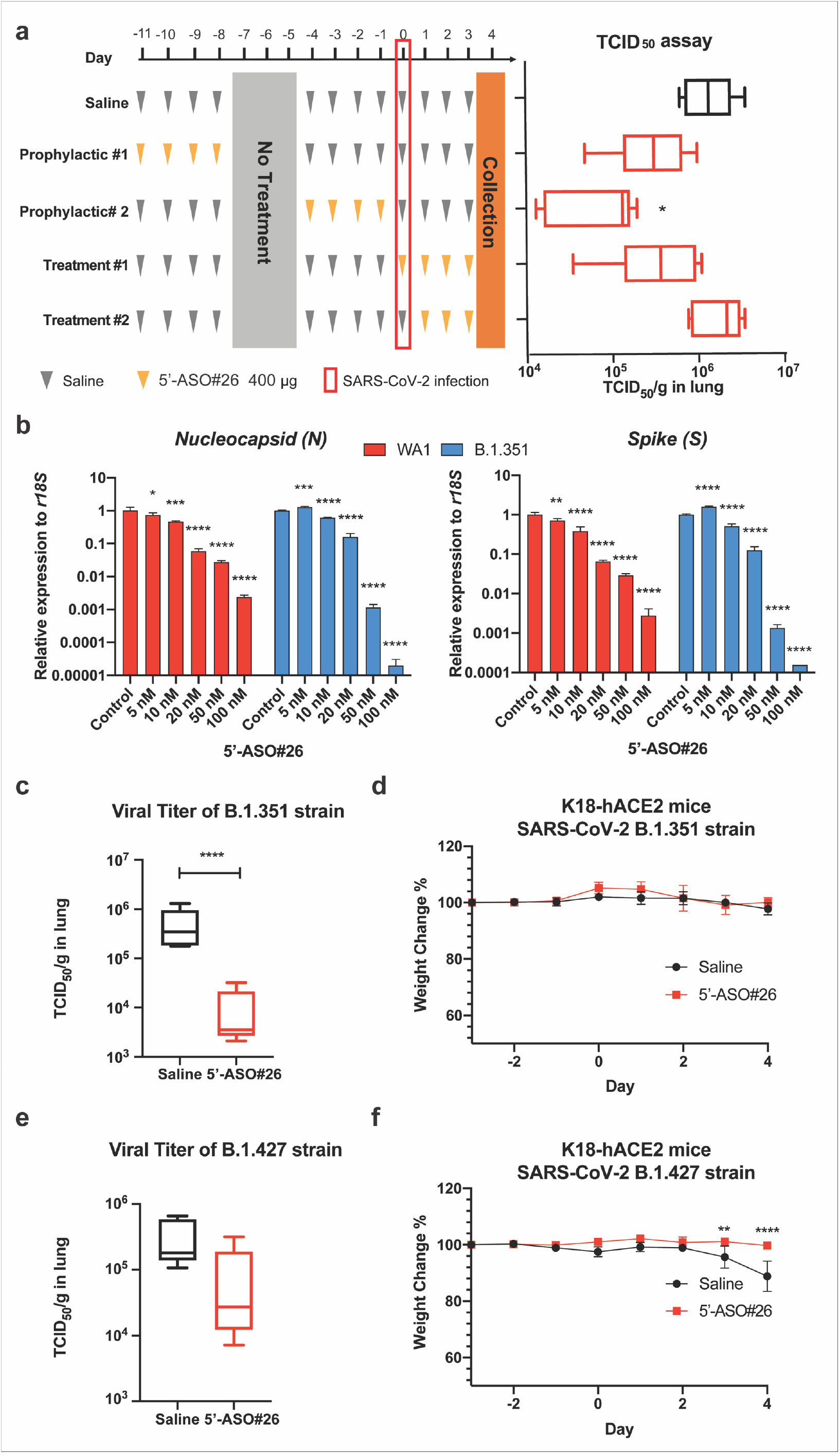
Prophylaxis and treatment of SARS-CoV-2 strains with 5’-ASO#26. a) Mice were administered with different treatment regimens of 5’-ASO#26 as indicated. Treatment on the day of infection was carried out at 6 hpi. The viral burden in lungs of mice in each group (N =5) was measured by TCID_50_ assay using lung homogenates. One-way ANOVA with Dunnett’s test was used to determine significance (* *P* < 0.05). b) Dose-dependent effects of 5’-ASO#26 on inhibition of viral replication of SARS-CoV-2 WA1 and B.1.351 strains were evaluated in infected Huh-7 cells by RT-qPCR of Nucleocapsid protein (N) and Spike (S) RNA. One-way ANOVA with Dunnett’s test was used to determine significance (** *P* < 0.01, **** *P* < 0.0001). c) and e) The viral burden in lungs of mice treated with Saline (N=5) and 5’-ASO#26 (N =5) was measured by TCID_50_ assay using lung homogenates. Student *t*-test was used to determine significance (**** *P* < 0.0001). d) and f) Weight change was monitored (N=5, symbols represent mean ± SD). Two-way ANOVA was used to determine significance (** *P* < 0.01, **** *P* < 0.0001). For a), c) and e), center line, median; box limits, upper and lower quartiles; plot limits, maximum and minimum in the boxplot.

Because the 5’ leader sequence of SARS-CoV-2 is highly conserved and as 5’-ASO#26 does not target Spike, we predicted that 5’-ASO#26 should also be able to repress the replication of SARS-CoV-2 variant strains. Therefore, we tested several reported variants with 5’-ASO#26 in cell-based assays. Our results showed that regardless of the mutations, 5’-ASO#26 exhibits potent repressive activity on viral replication of multiple SARS-CoV-2 variants, including B.1.351, D614G, B.1.1.7, and B.1.427 (Fig. 4b and Extended Data Fig. 4b). We further tested the effect of 5’-ASO#26 *in vivo* to repress the B.1.427 and B.1.1.7 strains in K18-hACE2 mice following the same schedule showed in Fig. 3a. We performed both TCID_50_ assays and RT-qPCR of lung homogenates to assess viral titer and observed that 5’-ASO#26 can also block the viral replication of both variant strains *in vivo* (Fig. 4c, 4e and Extended Data Fig. 4c). Although mice were inoculated with same amount of variant strains, the *in vivo* replication rate of B. 1.427 was significantly lower than that of B. 1.351, however the viral titers of both variant strains were repressed to a similar degree by LNA ASO treatment (Fig. 4c and 4e). Of note, similar to animals infected with the WA1 strain, the B.1.351 strain did not induce weight loss at 4 dpi (Fig. 4d). However, we noticed that there was a significant weight loss induced by B.1.427 strain infection from 3 dpi in the saline-treated group (Fig. 4f). In contrast, the LNA ASO-treated mice did not exhibit significant weight loss after B.1.427 strain infection (Fig. 4f), indicating that the viral repressive effect of LNA ASO treatment is still able to prevent the progression of SARS-CoV-2-induced pathologies.

Antisense therapy is currently used in clinical treatment for a range of different diseases, including cytomegalovirus retinitis (Fomivirsen^31^), Duchenne muscular dystrophy (Eteplirsen^32^), and Spinal Muscular Atrophy (Nusinersen)^33^. Here we show for the first time that intranasally administered LNA ASO treatment represents a promising therapeutic strategy for virus-induced respiratory diseases such as COVID-19. Our study demonstrated that naked LNA ASOs delivered intranasally in saline exhibit potent efficacy *in vivo*, and no formulation is necessary to achieve therapeutic effect. Considering the relatively small-sized genomes of RNA viruses and the ability to rapidly determine the sequence of any viral genome by next-generation sequencing (NGS) techniques, the design and screening of anti-viral LNA ASOs can be very fast and efficient, allowing for a rapid response to other global health crises posed by emerging viral threats in the future. The apparent ability of single-stranded RNA viruses to accumulate immune-evading mutations presents great challenges for vaccine and therapeutics development. Of note, LNA ASOs can overcome the challenge of mutations due to the ability to design sequences specifically targeted to highly conserved and critical regulatory regions of the viral genome. Additionally, LNA ASO cocktails targeting multiple essential genomic regions of viruses may further increase the efficacy of LNA ASOs as therapeutic candidates to overcome viral evasion mutations. In conclusion, we have identified an intranasally delivered LNA ASO targeting the 5’ leader sequence as a viable therapeutic approach for preventing or treating SARS-CoV-2 infections, including those caused by variants of concern, indicating that LNA ASOs can be pursued as lead candidates for the treatment of COVID-19.

## Methods

### Cell culture and viruses

Huh7 and Vero E6 cells were cultured in Dulbecco’s Modified Eagle Medium (DMEM) supplemented with 10% fetal bovine serum (FBS) and 1% Penicillin/Streptomycin. Cells were maintained at 37°C in 5% CO_2_. Transfection in Huh7 cells was performed with Lipofectamine 3000 reagent (Thermo Fisher). Cells were infected by SARS-CoV-2 viruses 12 hrs after transfection. The 2019n-CoV/USA_WA1/2020 isolate of SARS-CoV-2 was obtained from the US Centers for Disease Control and Prevention. Infectious stocks were produced by inoculating Vero E6 cells and collecting the cell culture media upon observation of cytopathic effect; debris were removed by centrifugation and passage through a 0.22 μm filter. The supernatant was then aliquoted and stored at −80°C. D614G, B.1.427 and B.1.1.7 strains were kind gifts from Dr. Mary Kate Morris at the California Department of Public Health (CDPH).

### Locked nucleic acid antisense oligonucleotides (LNA ASOs)

LNA ASOs were purchased from Integrated DNA Technologies (IDT). For the screening, small scale synthesis (100 nmole) of all LNA ASOs was followed by standard desalting purification. For the animal trials, large scale synthesis (250 mg) of the 5’-ASO#26 LNA ASO was followed by HPLC purification and endotoxin test.

### Biosafety

All aspects of this study were approved by the office of Environmental Health and Safety at UC Berkeley before initiation of this study. Work with SARS-CoV-2 was performed in a biosafety level 3 laboratory by personnel equipped with powered air-purifying respirators.

### Animals

C57BL/6J and K18-hACE2 [B6.Cg-Tg(K18-ACE2)2Prlmn/J] mice were purchased from the Jackson Laboratory and male golden Syrian hamsters at 4–5 weeks old were obtained from Charles River Labs (Strain Code:049). K18-hACE2 mice or hamsters were anesthetized using isoflurane and inoculated with 1 × 10^4^ TCID_50_ (for mice) or 10 TCID_50_ (for hamsters) of SARS-CoV-2 intranasally. For the treatment, saline (40 μl) or 5’-ASO#26 (indicated amount of LNA ASO in 40 μl saline) was administered once-daily intranasally. Mice and hamsters were sacrificed at 4 days post infection (dpi). The left lung lobe was collected and lysed for fifty-percent tissue culture infective dose (TCID_50_) assay, the inferior lobe was collected for RNA extraction and the post-caval lobe was collected for histological analysis. All procedures involving the use of mice and hamsters were approved by the University of California, Berkeley Institutional Animal Care and Use Committee. All protocols conform to federal regulations, the National Research Council Guide for the Care and Use of Laboratory Animals, and the Public Health Service Policy on Humane Care and Use of Laboratory Animals.

### Fifty-percent tissue culture infective dose (TCID_50_) assay

Virus viability and titers were evaluated in TCID_50_ assay within Vero E6 cells. Briefly, ten thousand cells were plated in each well in 96-well plates and cultured at 37°C overnight. Medium from SARS-CoV-2-infected cells or lysates from mouse lungs were used for ten-fold serial dilution with DMEM and added to the 96-well plates of Vero E6 cells. The plates were observed for cytopathic effect (CPE) after 3 days of culturing. The TCID_50_ results were calculated using the Spearman and Karber method^34^.

### RNA extraction and real-time quantitative PCR (RT-qPCR)

Infected Huh-7 cells (with or without medium) or mouse lung tissues were lysed in DNA/RNA shield reagent (Zymo Research) and total RNA was extracted by using RNeasy kit (Qiagen) according to the manufacturer’s protocol. cDNA was prepared by iScript™ Reverse Transcription Supermix (BioRAD) and qPCR was performed with Fast SYBR™ Green Master Mix (Thermo Fisher) and the reaction was run on the QuantStudio6 System (Applied Biosystems). mRNA levels were normalized to that of r18S. qPCR primer sets are as follow: SARS-CoV-2 Nucleocapsid(N): Fw 5’-GACCCCAAAATCAGCGAA AT-3’ and Rv 5’-TCTGGTTACTGCCAGTTGAATCTG-3’, SARS-CoV-2 Spike(S): Fw 5’-GTCCTTCCCTCAGTCAGCAC-3’ and Rv 5’-ATGGCAGGAGCAGTTGTGAA-3’, Human r18S: Fw 5’-GTAACCCGTTGAACCCCATT-3’ and Rv 5’-CCATCCAATCGGTAGTAGCG-3’, Mouse r18S: Fw 5’-GCAATTATTCCCCATGAACG -3’ and Rv 5’ - GGCCTCACTAAACCATCCAA -3’.

### RNA-sequencing

cDNA libraries were constructed from 500 ng of total RNA from Huh-7 or lung tissues of mice according to the manufacturer’s protocol of Stranded mRNA-seq kit (KAPA). Briefly, mRNA was captured and fragmentized to 100~300 bp. library construction was performed undergoing end repair, A tailing, ligation of unique dual-indexed adapters (KAPA) and amplification of 10 cycles to incorporate unique dual index sequences. Libraries were sequenced on the NovaSeq 6000 (Novogene) targeting 40 million read pairs and extending 150 cycles with paired end reads. The data have been deposited in NCBI’s Gene Expression Omnibus and are accessible through GEO Series accession number GSE174382. The published RNA-seq data of infected K18-hACE2 at 0 dpi and 4 dpi were obtained from GSE154104. STAR aligner^35^ was used to map sequencing reads to transcripts in the mouse mm10 reference genome. Read counts for individual transcripts were produced with HTSeq-count^36^, followed by the estimation of expression values and detection of differentially expressed transcripts using EdgeR^37^. Differentially expressed genes were defined by at least 2-fold change with FDR less than 0.01.

### DMS modification of *in vitro*-transcribed RNA

gBlock containing the first 3,000 nucleotides of the SARS-CoV-2 genome (2019-nCoV/USA-WA1/2020) was obtained from IDT. The gBlock was amplified by PCR with a forward primer that contained the T7 promoter sequence (5’-TAATACGACTCACTATAGGGATTAAAGGTTTATACCTTCCCAGGTAAC-3’) and the reverse primer (5’-TCGTTGAAACCAGGGACAAG-3’). The PCR product was used as the template for T7 Megascript in vitro transcription (ThermoFisher Scientific) according to manufacturer’s instructions. Next, 1 μl of Turbo DNase I (ThermoFisher Scientific) was added and incubated at 37°C for 15 min. The RNA was purified using RNA Clean and Concentrator™-5 (Zymo). Between 1-2 μg RNA was denatured at 95°C for 1 min. Denatured RNA was refolded in the presence of 2 μM of LNA ASO by incubating the mixture in 340 mM sodium cacodylate buffer (Electron Microscopy Sciences) and 5 mM MgCl^2+^, such that the volume was 97.5 μl, for 20 min at 37°C. Then, 2.5% DMS (Millipore-Sigma) was added and incubated for 5 min at 37°C whole shaking at 800 r.p.m. on a thermomixer. Subsequently, DMS was neutralized by adding 60 μl β-mercaptoethanol (Millipore-Sigma). The RNA was precipitated by incubating in 3 μl (45 μg) glycoblue coprecipitant (Invitrogen), 18 μl 3M sodium acetate and 700 μl ethanol between 1 h and overnight at −80°C, followed by centrifugation at max speed for 45 min in 4°C. The RNA was washed with 700 μl ice cold 75% ethanol and centrifuged for 5 min. RNA was resuspended in 10 μl water.

### DMS modification of infected Huh-7 cells with ASO treatment

Huh-7 cells were transfected with LNA ASO (50 nM) 12 h before the infection. After infection of SARS-CoV-2 (MOI 0.05), cells were cultured in 6-well plates with 2 ml of media. Then, 2.5% DMS was added to cells and incubated for 3 min at 37°C. Subsequently, after careful removal of the media, DMS was neutralized by adding 20 ml of chilled 10% β-mercaptoethanol in PBS. The cell pellets were washed once with chilled PBS and collected for RNA extraction.

### rRNA subtraction of total cellular RNA from DMS-treated cells

Between 3-5 μg RNA per sample was used as the input for rRNA subtraction. First, equal amount of rRNA pooled oligonucleotides were added and incubated in hybridization buffer (200 mM NaCl, 100 mM Tris-HCl, pH 7.4) in a final volume of 60 μl. The samples were denatured for 2 min at 95°C, followed by a reduction of 0.1°C/s until the reaction reached 45°C. 3-5 μl Hybridase™ Thermostable RNase H (Lucigen) and 7 μl 10x RNase H buffer preheated to 45°C was added. The samples were incubated at 45°C for 30 min. The RNA was purified using RNA Clean and Concentrator™-5 kit and eluted in 42 μl water. Then, 5 μl Turbo DNase buffer and 3 μl Turbo DNase (ThermoFisher Scientific) were added to each sample and incubated for 30 min at 37°C. The RNA was purified using RNA Clean and Concentrator™-5 kit and eluted in 10 μl water.

### RT-PCR and sequencing of DMS-modified RNA

To reverse transcribe, rRNA-depleted total RNA or in vitro-transcribed RNA purified from the previous steps was added to 4 μl 5x FS buffer, 1 μl dNTP, 1 μl of 0.1 M DTT, 1 μl RNase Out, 1 μl of 10 μM reverse primer (5’-TCGTTGAAACCAGGGACAAG-3’) and 1 μl TGIRT-III (Ingex). The reaction was incubated for 1.5 h at 60°C. Then, to degrade the RNA, 1 μl of 4 M NaOH was added and incubated for 3 min at 95°C. The cDNA was purified in 10 μl water using the Oligo Clean and Concentrator™ kit (Zymo). Next, 1 μl of cDNA was amplified using Advantage HF 2 DNA polymerase (Takara) for 25-30 cycles according to the manufacturer’s instructions (Fw 5’-GGGATTAAAGGTTTATACCTTCCC-3’ and Rv 5’-TCGTTGAAACCAGGGACAAG-3’). The PCR product was purified using E-Gel™ SizeSelect™ II 2% agarose gel (Invitrogen). RNA-seq library for 300 bp insert size was constructed following the manufacturer’s instructions (NEBNext Ultra™ II DNA Library Prep Kit). The library was loaded on iSeq-100 Sequencing flow cell with iSeq-100 High-throughput sequencing kit and library was run on iSeq-100 (paired-end run, 151 x 151 cycles).

### Immunohistochemistry (IHC) and hematoxylin and eosin (H&E) staining

Histology was performed by HistoWiz Inc. (histowiz.com) using a Standard Operating Procedure and fully automated workflow. Samples were processed, embedded in paraffin, and sectioned at 4 μm. Immunohistochemistry was performed on a Bond Rx autostainer (Leica Biosystems) with enzyme treatment (1:1000) using standard protocols. Antibodies used were rabbit monoclonal CD3 primary antibody (Abcam, ab16669, 1:100), rabbit monoclonal B220 primary antibody (Novus, NB100-77420, 1:10000), rabbit monoclonal SARS-CoV-2 (COVID-19) nucleocapsid primary antibody (GeneTex, GTX635686, 1:8000) and rabbit anti-rat secondary (Vector, 1:100). Bond Polymer Refine Detection (Leica Biosystems) was used according to the manufacturer’s protocol. After staining, sections were dehydrated and film coverslipped using a TissueTek-Prisma and Coverslipper (Sakura). Whole slide scanning (40x) was performed on an Aperio AT2 (Leica Biosystems).

### Statistical analysis

Data are presented as mean values, and error bars represent SD. Data analysis was performed using GraphPad Prism 8. Data were analyzed using unpaired t-test; one-way or two-way ANOVA followed by Turkey or Dunnett test as indicated. P value < 0.05 was considered as statistically significant.

### Data availability

All data are available in the manuscript and associated files. Source data is provided with this paper.

## Acknowledgements

We thank Dr. Fyodor Urnov and Dr. Jennifer Doudna and the Innovative Genomics Institute for organizing the COVID-19 research efforts at UC Berkeley and orchestrating financial support, as well as Dr. Hesong Han, Dr. Jie Li, Dr. Aaron Mendez, Dr. Niren Murthy, Dr. Britt Glaunsinger and Angelica M. Gonzalez at UC Berkeley for experimental design advice and technical assistance. The D614G, B.1.427 and B.1.1.7 SARS-CoV-2 strains were kind gifts from Dr. Mary Kate Morris at California Department of Public Health (CDPH). The research was supported by the following funding sources: Fast Grants (Emergent Ventures at the Mercatus Center, George Mason University) to A.M.N., S.S., and E.H., Innovative Genomics Institute grant (A.M.N.), and UC Berkeley/Anonymous Donor (A.M.N.).

## Author contributions

C.Z. and A.M.N. designed and organized the study, reviewed all data; C.Z., J.Y.L. and A.M.N. wrote and all authors reviewed and edited the manuscript; S.K. and A.M.N. designed the LNA ASOs; C.Z., L.H.Y., F.G., D.F., and S.S. carried out and/or supervised the animal experiments in BSL-3; C.Z., X.N., S.B.B., E.V.D., E.H. and S.S. carried out and/or supervised *in vitro* experiments in BSL-3; C.Z., J.Y.L. and L.X. conducted the BSL-2 work and NGS sample preparation; J.Z.W. and S.R. carried out the DMS-seq and analysis; F.J. and R.I.S. did the RNA-seq analyses; C.C. isolated and provided the B.1.427 variant, A.R., B.A.P. and C.A.B. isolated and provided the B.1.351 variant.

## Figures and Legends

**Extended Data Figure 1.**
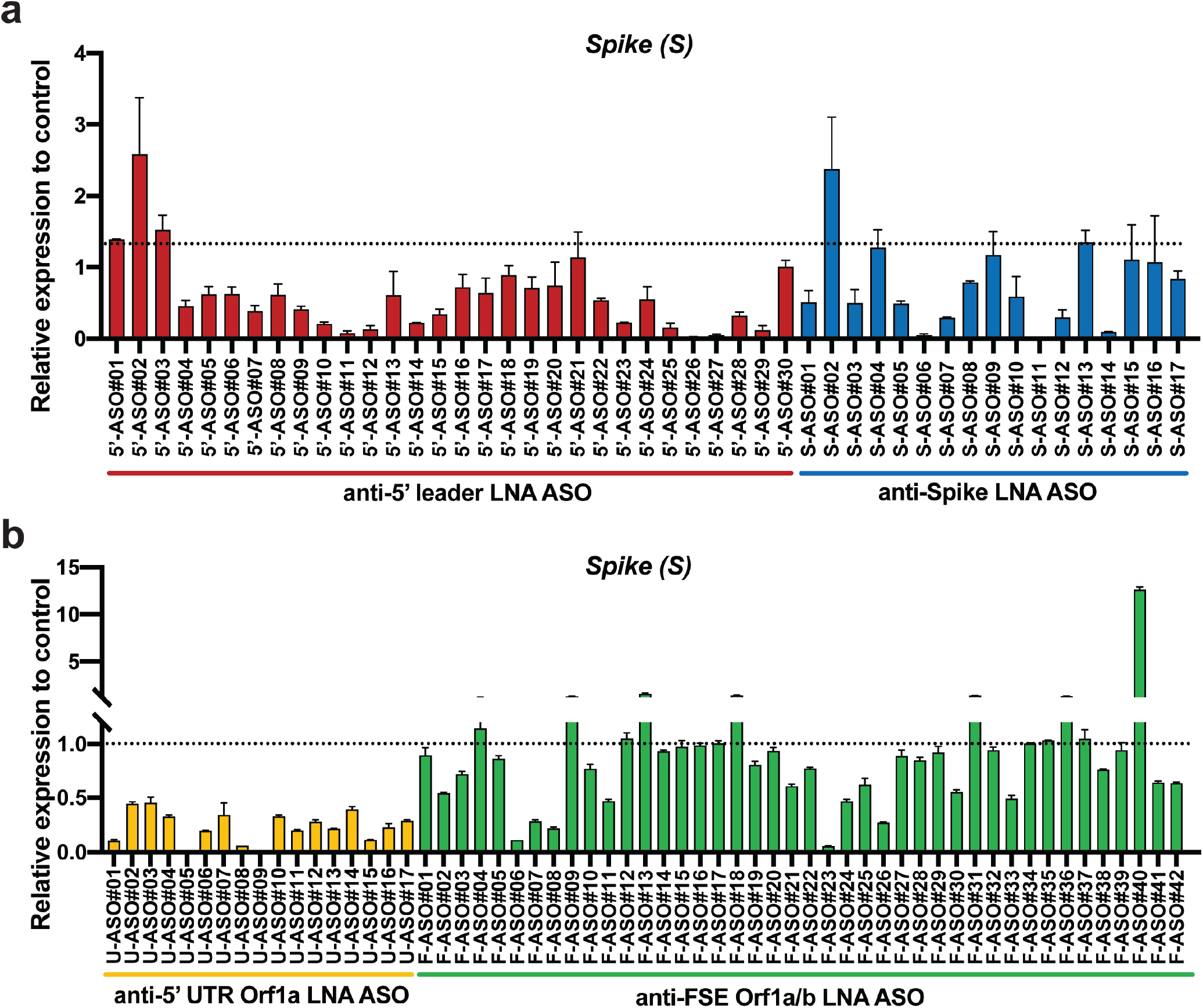
*In vitro* screening of LNA ASOs targeting SARS-CoV-2. a) and b) Infected Huh-7 cells were treated with LNA ASO (100 nM) and cell culture media was collected at 48 hpi. Levels of viral Spike (S) RNA were analyzed by RT-qPCR. Each LNA ASO was tested in duplicate. One-way ANOVA with Dunnett’s test was used to determine significance (**** *P* < 0.0001).

**Extended Data Figure 2.**
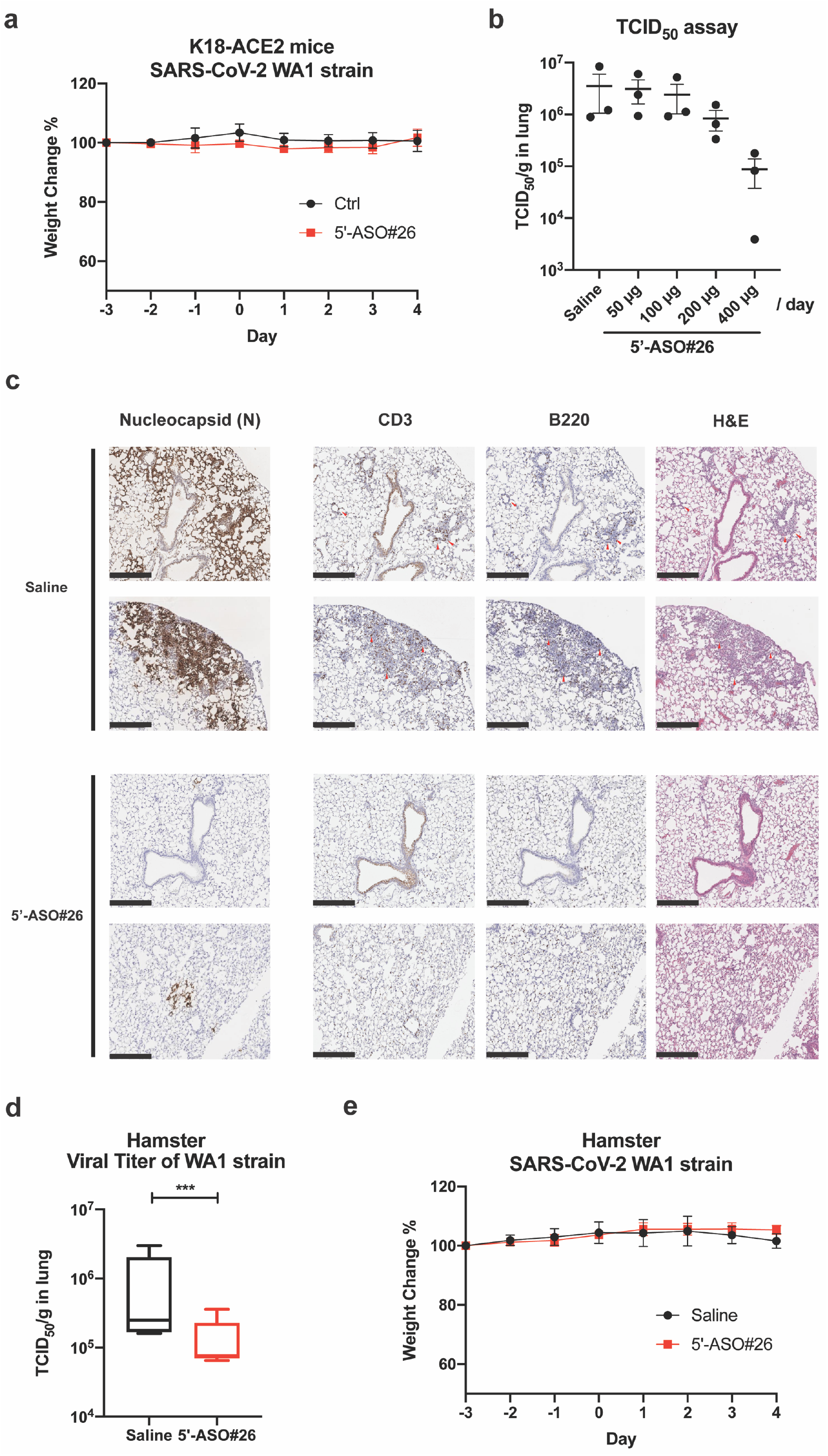
Physiological and histological analysis of LNA ASO-treated K18-hACE2 mice. a) Weight change of mice was monitored over the treatment/infection course (N=5 in each group, symbols represent mean ± SD). b) LNA ASO dose-dependence efficacy testing in mice. Mice were administered with different doses of 5’-ASO#26 in 40 μl saline as indicated. The viral burden in lungs of mice in each group (N=3) were measured by TCID_50_ assay using lung homogenates. c) Representative images of hematoxylin and eosin (H&E) staining of lung sections and immunohistochemistry (IHC) staining of CD3 and B220 in infected K18-hACE2 mice with Saline (N=5) or 5’-ASO#26 treatment (N=5). Scale bar = 500 μm. d) The viral burden in lungs of hamsters treated with Saline (N=5) or 5’-ASO#26 (N=5) was measured by TCID_50_ assay using lung homogenates. Center line, median; box limits, upper and lower quartiles; plot limits, maximum and minimum in the boxplot. e) Weight change of hamsters was monitored over the treatment/infection course (N=5 in each group, symbols represent mean ± SD). For d) Student *t*-test was used to determine significance (*** *P* < 0.001) and for a) and e), Two-way ANOVA was used to determine significance.

**Extended Data Figure 3.**
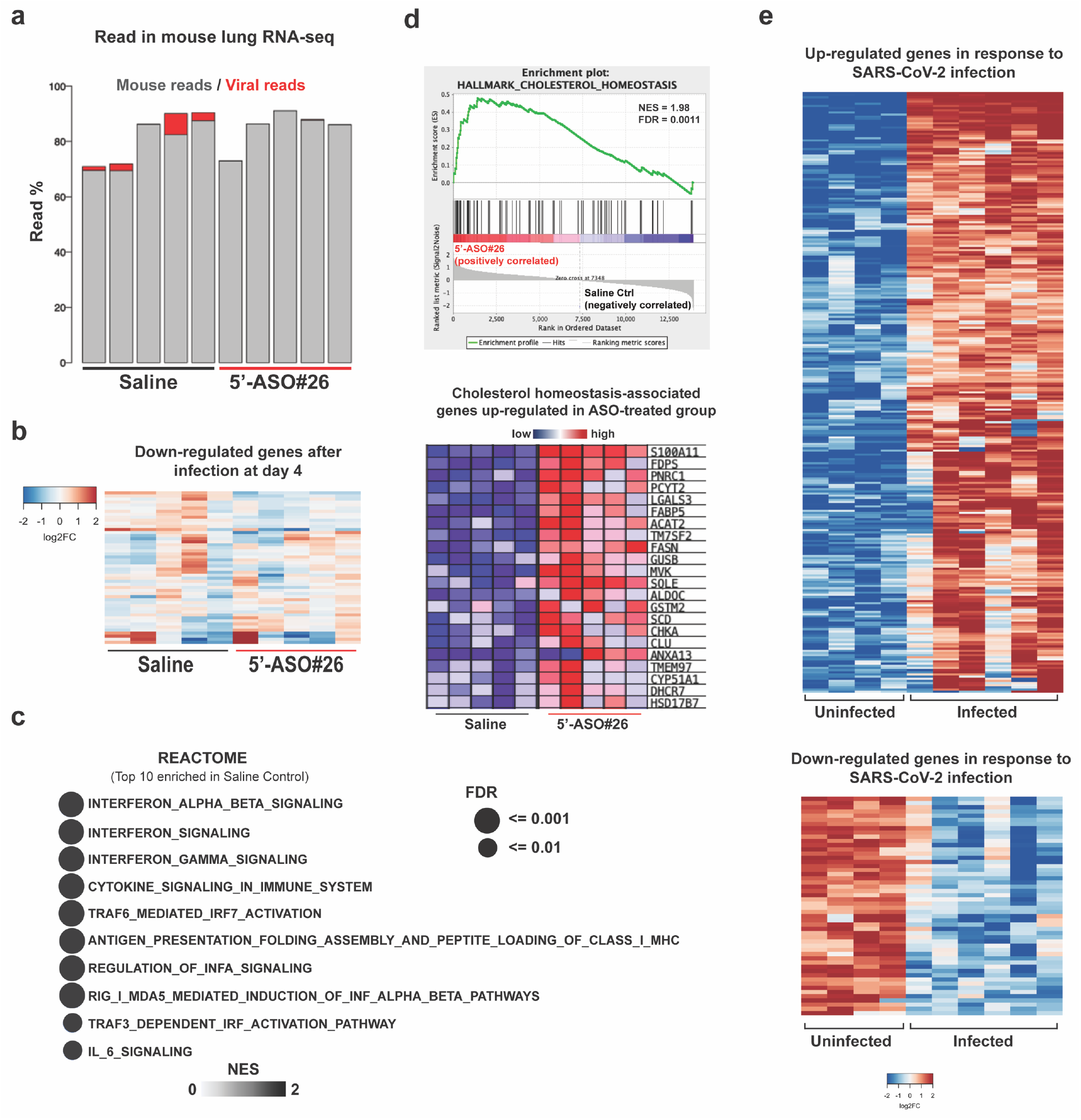
Evaluating the *in vivo* effects of 5’-ASO#26 in K18-hACE2 mice. a) Percentage of human/viral sequencing reads in RNA-seq data of Saline- and LNA ASO-treated mouse lung. Reads mapped to the human genome marked in gray and reads mapped to virus marked in red. b) Expression change of SARS-CoV-2 infection-downregulated genes in Saline- and LNA ASO-treated group. Infection. c) GSEA of REACTOME gene sets enriched among upregulated genes in lungs of Saline-treated mice. Terms were ranked by the false discovery rate (q value). d) GSEA plot and heatmap of significantly upregulated genes enriched in cholesterol homeostasis pathway in ASO-treated mice. e) Heatmaps of significantly upregulated and downregulated (> 2-fold, FDR<0.01) genes at day 4 of SARS-CoV-2 infection. For b), d) and e) Columns represent samples and rows represent genes. Colors indicate gene expression levels (log2 RPKM) relative to average expression across all samples.

**Extended Data Figure 4.**
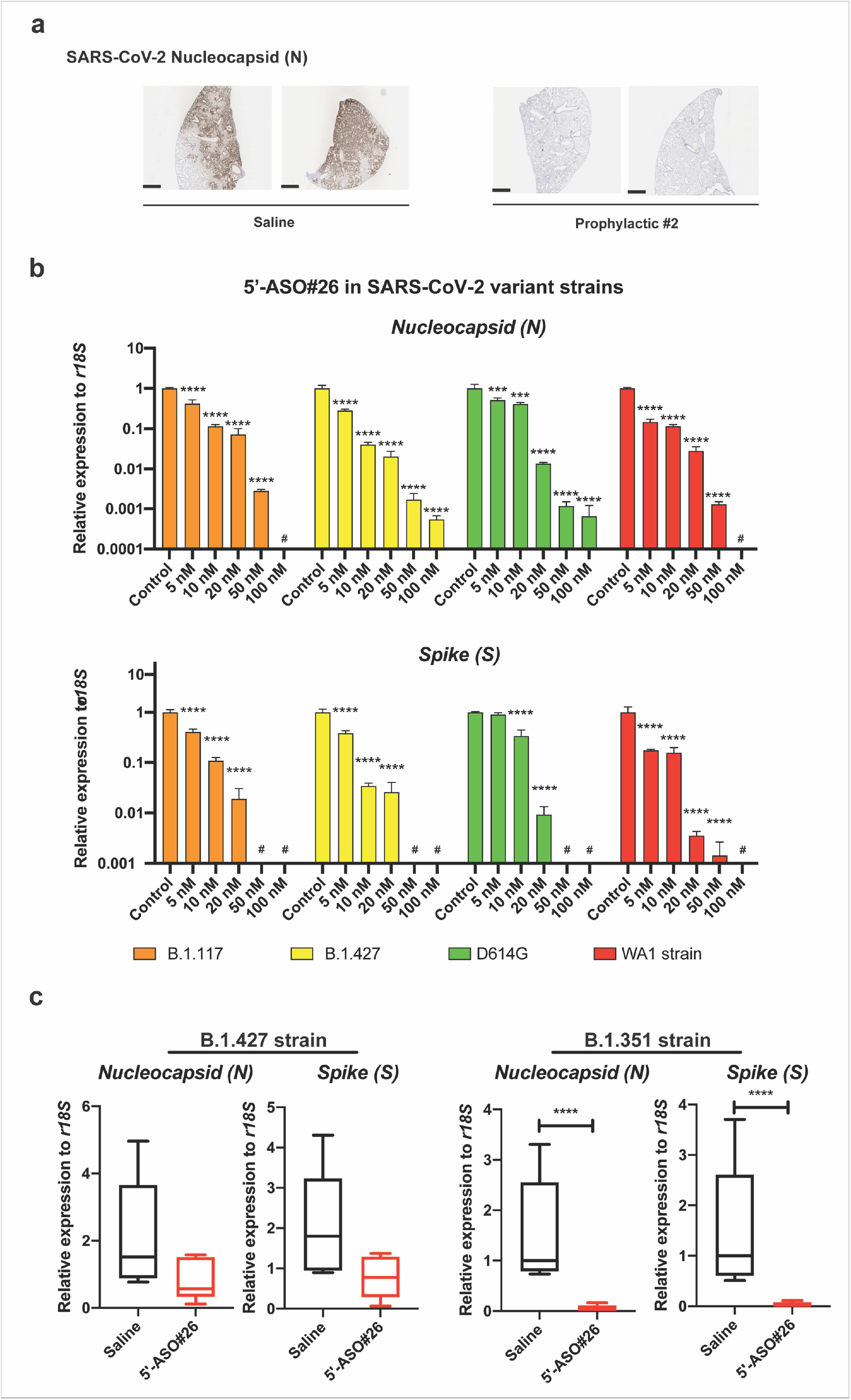
5’-ASO#26 is capable of repressing the replication of SARS-CoV-2 variant strains. a) Representative images of IHC staining of SARS-CoV-2 nucleocapsid protein in Saline (N=5) or Prophylactic#2 (N=5) regimen groups. Scale bar = 2 mm. b) Dose-dependent efficacy of 5’-ASO#26 in repressing replication of SARS-CoV-2 variant strains was evaluated in infected Huh-7 cells with increasing doses of 5’-ASO#26 by RT-qPCR of Nucleocapsid (N) and Spike (S) RNAs. One-way ANOVA with Dunnett’s test was used to determine significance (*** *P* < 0.001, **** *P* < 0.0001, # no detection). c) Viral RNA levels in mouse lungs were analyzed by RT-qPCR of Nucleocapsid (N) and Spike (S) RNAs. Center line, median; box limits, upper and lower quartiles; plot limits, maximum and minimum in the boxplot. Student *t*-test was used to determine significance (* *P* < 0.05, **** *P* < 0.0001).

**Extended Data Table 1.**
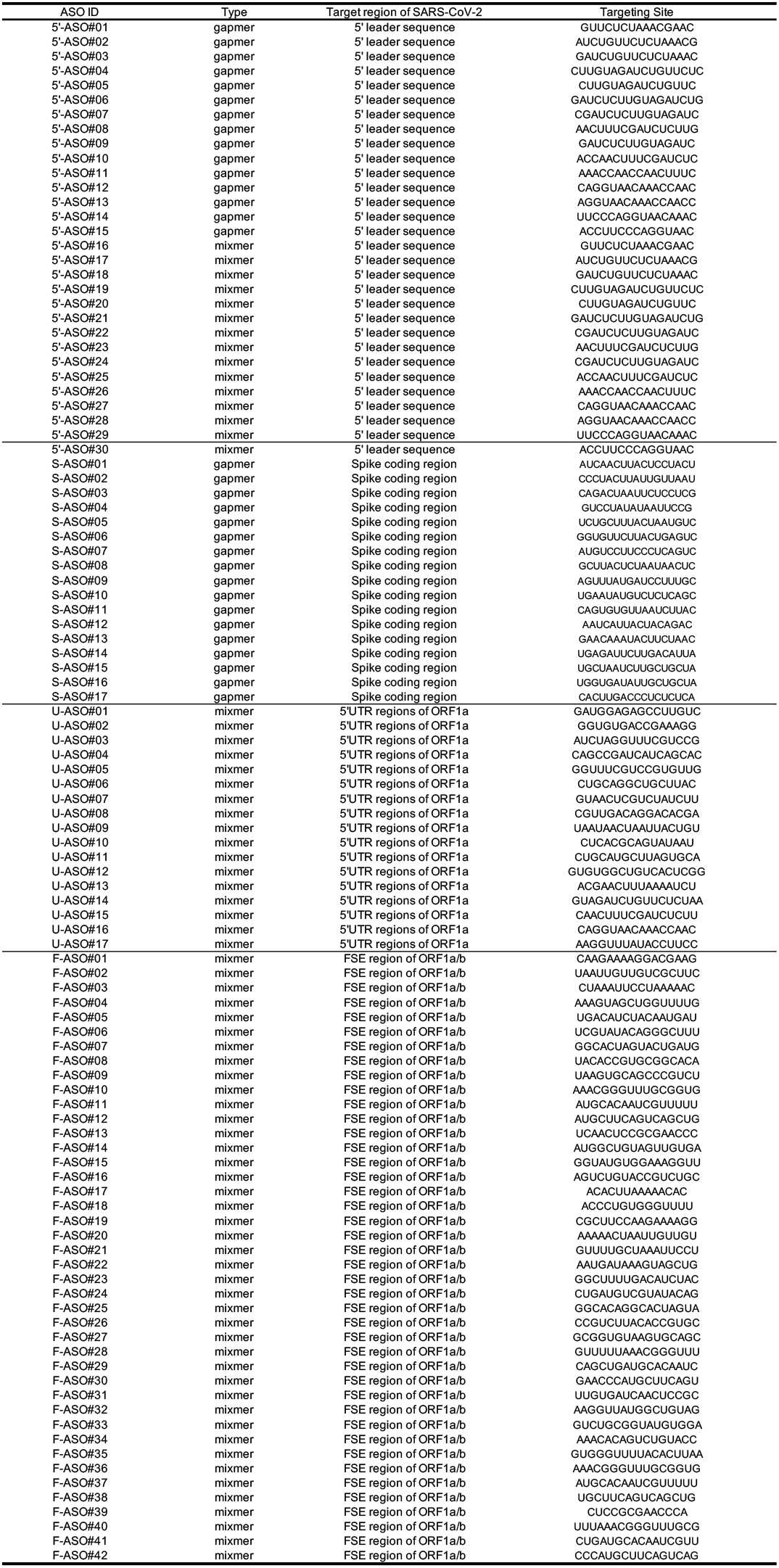
Targeting sites of LNA ASOs used in *in vitro* screening. To identify LNA ASOs exhibiting anti-viral efficacy, LNA ASOs targeting 5’ leader sequences, 5’ UTR region of ORF1a, FSE of ORF1a/b and Spike coding region were tested in cell-based screening assays in Huh-7 cells.

